# Suppression of DSB Formation by Polβ in Active DNA Demethylation is Required for Postnatal Hippocampal Development

**DOI:** 10.1101/852053

**Authors:** Akiko Uyeda, Kohei Onishi, Teruyoshi Hirayama, Satoko Hattori, Tsuyoshi Miyakawa, Takeshi Yagi, Nobuhiko Yamamoto, Noriyuki Sugo

## Abstract

Genome stability is essential for brain development and function. However, the contribution of DNA repair to genome stability in neurons remains elusive. Here, we demonstrate that the base excision repair protein Polβ is involved in hippocampal neuronal differentiation via a TET-mediated active DNA demethylation during early postnatal stages. Polβ deficiency induced extensive DNA double-strand breaks (DSBs) in hippocampal neurons, and a lesser extent in cortical neurons, during a period in which decreased levels of 5-methylcytosine were observed in genomic DNA. Inhibition of the hydroxylation of 5-methylcytosine by microRNAs miR29a/b-1 expression diminished DSB formation. Conversely, its induction by TET1 overexpression increased DSBs. The damaged hippocampal neurons exhibited aberrant neuronal gene expression profiles and dendrite formation. Behavioral analyses revealed impaired spatial learning and memory in adulthood. Thus, Polβ maintains genome stability in the active DNA demethylation that occurs during postnatal neuronal development, thereby contributing to differentiation and subsequent behavior.

## Introduction

Genome stability is crucial for both genetic and epigenetic regulation underlying gene expression in the brain throughout life. DNA repair is essential to maintain genome stability and has been well characterized through studies on cancer and immune cell differentiation in mammals (Alt et al, 2013; Lindahl & Wood, 1999). In the nervous system, mouse models reveal that DNA repair dysfunction in neural progenitors frequently leads to genome instability and neuronal apoptosis during the period of neurogenesis (Deans et al, 2000; Gao et al, 1998; Gu et al, 2000; Lee et al, 2001; Lee et al, 2009; Pulvers & Huttner, 2009; Sugo et al, 2000). In addition, genetic diseases related to DNA repair defects include microcephaly, developmental disorders, and psychiatric disorders (Madabhushi et al, 2014; McKinnon, 2013). Accumulation of somatic mutations in neurons during development has been implicated in developmental brain disorders such as autism and schizophrenia (McConnell et al, 2017; McKinnon, 2013; Poduri et al, 2013). These studies suggest that DNA repair is likely to be critical for normal brain development and function. However, while DNA repair has been characterized in mitotic cells including neural progenitors, its role in neurons as postmitotic cells remains unclear. In brain development, postnatal neuronal differentiation is also a core process for circuit formation and activity-dependent refinement (Flavell & Greenberg, 2008; Kolodkin & Tessier-Lavigne, 2011). Thus, it is important to uncover novel aspects of DNA repair in neuronal differentiation and function.

Base excision repair (BER) is mainly involved in the removal of DNA base damage and apurinic/apyrimidinic sites (Wilson et al, 2000). In addition, recent studies have revealed that BER also plays a role in the active DNA demethylation process as an epigenetic regulation (Schuermann et al, 2016; Wu & Zhang, 2010). In this process, 5-methylcytosine (5mC) is initially oxidized by TET enzymes and is converted to 5-hydroxymethylcytosine (5hmC) (Ito et al, 2010; Ito et al, 2011; Tahiliani et al, 2009); the modified base is finally recognized by thymine DNA glycosylase and replaced with cytosine by DNA polymerase β (Polβ) and the Xrcc1/Lig3 complex (Cortellino et al, 2011, Cortázar et al, 2011, He et al, 2011, Weber et al, 2016). DNA methylation and demethylation often play a central role in cell differentiation (Moore et al, 2013; Wu & Zhang, 2017). In the neuronal epigenome, dynamic changes in the DNA methylation level are observed during brain development (Lister, 2013; Sharma, 2016; Simmons et al, 2013) and affect neuronal gene expression, which is implicated in neurogenesis, maturation, and plasticity (Feng et al, 2010; Moretti, et al, 2006; Sanosaka et al, 2017). This regulation also contributes to learning and memory (Gontier et al, 2018; Kaas et al, 2013; Li et al, 2014; Rudenko et al, 2013).

Studies using conventional *Polβ*-deficient mice show increased neuronal apoptosis during the period of neurogenesis in the developing nervous system rather than in other tissues, and the mice die just after birth (Sugo et al, 2000). The p53-dependent pathway regulates neuronal apoptosis after the final mitosis (Sugo et al, 2007; Sugo et al, 2004). Our previous study focusing on spatiotemporal roles using forebrain-specific conditional knockout *Emx1-Cre/Polβ^fl/fl^* and *Nex-Cre/Polβ^fl/fl^* mice indicates that Polβ deficiency in neural progenitors rather than in postmitotic neurons specifically leads to an increase of DNA double-strand breaks (DSBs) associated with replication in the embryonic cortex (Onishi et al, 2017). The accumulation of DSBs frequently induces neuronal apoptosis and abnormal axon projection. Furthermore, impairment of the DNA demethylation process is a potential cause of DSBs in Polβ-deficient progenitors, suggesting that epigenetic regulation via BER including Polβ in neural progenitors is essential for neuronal survival and differentiation. However, how Polβ contributes to subsequent neuronal development, gene expression, and further cognitive function is not fully understood.

To address this issue, we investigated the role of Polβ using *Nex-Cre/Polβ^fl/fl^* mice, in which postmitotic excitatory neurons lack Polβ expression. We found that the mutant mice exhibited extensive DSB formation, but not apoptosis, in hippocampal neurons more so than in cortical neurons during early postnatal stages, in which the levels of 5mC and 5hmC in the genome decreased. *In vivo* manipulation of active DNA demethylation during this period altered the extent of DSBs in Polβ-deficient neurons. Furthermore, Polβ deficiency affected gene expression profiles and dendritic morphology of developing hippocampal neurons, and impaired hippocampus-related learning and memory. These findings suggest that genome stability mediated by Polβ is required for active DNA demethylation leading to normal postnatal neuronal development and memory function.

## Results

### Polβ-deficient neurons show accumulation of DNA double-strand breaks in postnatal development

To investigate the spatiotemporal role of Polβ in postmitotic neuronal development, we used *Nex-Cre/Polβ^fl/fl^* mice (Onishi et al, 2017). In control *Polβ^fl/fl^* mice, Polβ immunoreactivity was roughly ubiquitous throughout the neocortex and the hippocampus at P2, and its subcellular localization was predominantly nuclear (Supplemental Figure (Figure S) 1A, B). As expected, excitatory neurons including Ctip2-positive cells lost Polβ expression in the neocortex and the hippocampus of *Nex-Cre*/*Polβ^fl/fl^* mice (Figure S1A, B). However, the cortical laminar organization and hippocampal cytoarchitecture in *Nex-Cre*/*Polβ^fl/fl^* mice seemed to be similar to those in control *Polβ^fl/fl^* mice (Figure S1A-C).

To examine whether Polβ deficiency affects genome stability in neuronal development, DSB formation was investigated at embryonic (E16.5 and 18.5) and postnatal stages (P2, 15, 28 and 90) by immunohistochemical analysis with an antibody against γH2AX, a DSB marker (Onishi et al, 2017, Rogakou et al, 1999, Rogakou et al, 1998). Strong signals of γH2AX foci were frequently found in *Nex-Cre/Polβ^fl/fl^* hippocampal pyramidal cell nuclei at P15, while only a few cells were focus-positive in controls (Figure 1A, B). Quantitative analyses showed that both the number of foci in a nucleus and the fraction of focus-positive cells were significantly larger in *Nex-Cre/Polβ^fl/fl^* pyramidal cells than in control *Polβ^fl/fl^* (Figure 1D, E). Consistent with these observations, immunostaining with 53BP1, a protein involved in non-homologous end joining (NHEJ), also showed focus formation in *Nex-Cre/Polβ^fl/fl^* mice (Figure S1D), strongly indicating that the foci were due to DSB formation (Schultz et al, 2000). The developmental time course further demonstrated that γH2AX foci were undetectable during the embryonic stages (Figure S2A) and appeared from P2 (Figure 2A). The signals just peaked at P15 and then decreased until the 3-month adult stage (Figure 2A, C, D). Similarly, a fraction of neocortical neurons in *Nex-Cre/Polβ^fl/fl^* mice also exhibited a transient increase of γH2AX focus formation with a comparable developmental time course (Figure 1A, C-E and 2B, E, F), although the increase was less marked than in hippocampal pyramidal neurons (Figure 1D, E; the fraction of γH2AX focus-positive cells at P15 was 79% in the hippocampus and 20% in the neocortex). These results indicate that Polβ deficiency transiently increases DSB formation during postnatal neuronal development, although the extent of DSBs differs between brain regions.

**Fig. 1.**
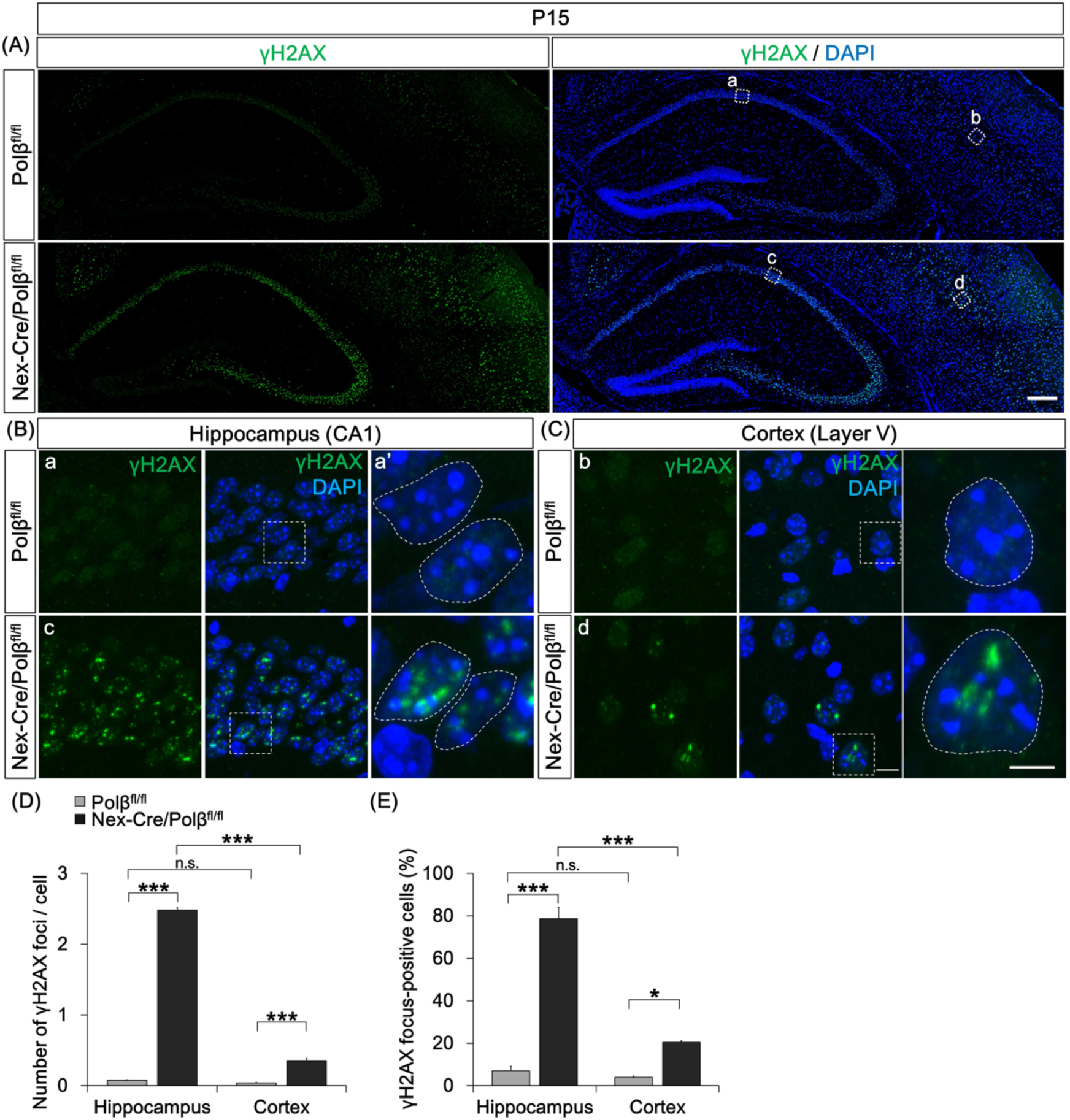
*Nex-Cre/Polβ^fl/fl^* mice exhibit DSBs in postnatal hippocampus and cortex. (A) Immunohistochemistry was performed with anti-γH2AX antibody in P15 *Nex-Cre/Polβ^fl/fl^* and *Polβ^fl/fl^* hippocampus and cortex. Scale bar, 400 µm. (B, C) Magnified images of the boxed areas in (A) including hippocampal CA1 pyramidal cell layer (a, c) and cortical layer V (b, d) are shown. Magnified images of the dashed-line boxed areas in the center image are shown in the the rightmost images. The dashed lines in the rightmost images mark the perimeter of the nucleus. Scale bars, 10 (the center) and 5 µm (the right). (D, E) Histograms show quantitative analysis of the mean number of γH2AX foci in each nucleus (D) and percentage of γH2AX foci-positive cells (E) in hippocampus and cortex of *Nex-Cre/Polβ^fl/fl^* (n = 492 cells, n = 666 cells) and *Polβ^fl/fl^* (n = 513 cells, n = 791 cells) mice. Data are the mean ± SEM from three different brains. Significant difference: *p < 0.05 and ***p < 0.001, ANOVA with Tukey’s post-hoc test.

**Fig. 2.**
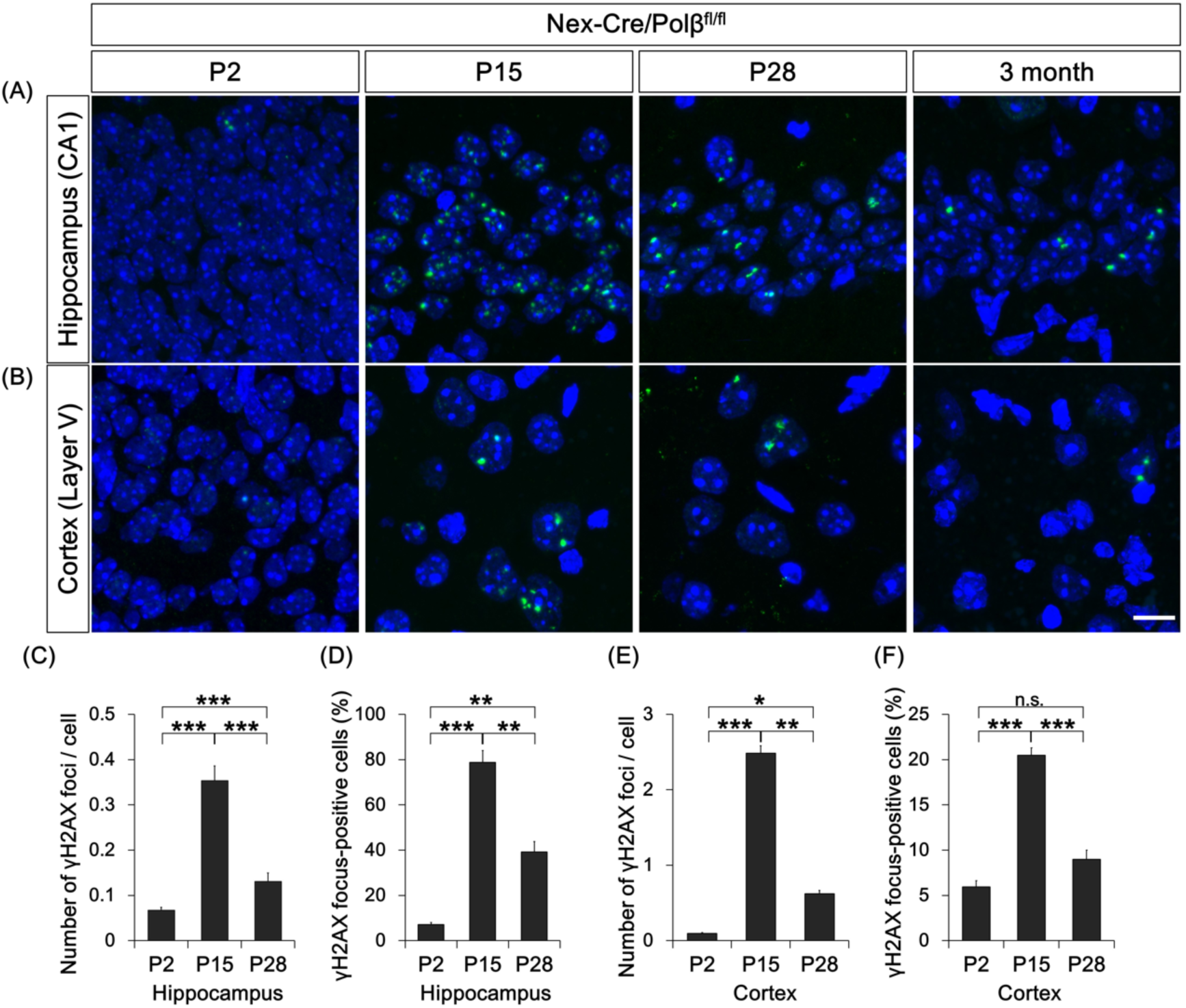
DSB formation in *Nex-Cre/Polβ^fl/fl^* mice transiently increases during postnatal development. (A, B) Immunohistochemistry was performed with anti-γH2AX antibody in *Nex-Cre/Polβ^fl/fl^* and *Polβ^fl/fl^* hippocampus (A) and cortex (B) at P2, P15, P28, and 3 months. Scale bar, 10 µm. (C-F) Histograms show quantitative analysis of the mean number of γH2AX foci in each nucleus (C, E) and γH2AX foci-positive cells (D, F) in hippocampus (C, D) and cortex (E, F) of *Nex-Cre/Polβ^fl/fl^* mice at P2 (n = 1160 cells, n = 2155 cells), P15 (n = 492 cells, n = 690 cells), and P28 (n = 477 cells, n = 620 cells). Data are the mean ± SEM from three independent experiments. Significant difference: *p < 0.05, **p < 0.01, and ***p < 0.001, ANOVA with Tukey’s post-hoc test.

A next question is whether DSB accumulation induces neuronal apoptosis as observed in *Emx1-Cre/Polβ^fl/fl^* mice (Onishi et al, 2017). Anti-cleaved caspase-3 immunohistochemistry was performed in *Nex-Cre/Polβ^fl/fl^* mice. Unexpectedly, few cleaved caspase 3-positive cells were observed in *Nex-Cre/Polβ^fl/fl^* hippocampus and cortex at P2 and P15, during which DSB formation increases. The abundance of apoptotic cells was similar to that in control *Polβ^fl/fl^* mice (Figure S2C). Taken together, these results suggest that Polβ deficiency leads to genome instability, but does not affect cell survival, in hippocampal and cortical neurons during postnatal development.

### Polβ is required for base excision repair in postmitotic neurons

Polβ is a key enzyme in BER but not in DSB repair (DSBR) (Sobol et al, 1996, Wilson et al, 2000). The DSB formation in Polβ-deficient neurons may be due to accumulation of single-strand breaks (SSBs) as BER intermediates (Caldecott, 2003). To test this possibility, immunohistochemical analysis with an antibody against XRCC1, an SSB marker (Caldecott, 2003, El-Khamisy et al, 2003), was performed. Fluorescence intensity of XRCC1 was significantly increased in P15 *Nex-Cre/Polβ^fl/fl^* hippocampal CA1 pyramidal cell nuclei compared to control *Polβ^fl/fl^* nuclei (Figure 3A-C). In addition, the XRCC1 intensity increased even during normal hippocampal development from P2 to P15 (Figure 3A, C), similar to developmental changes of the γH2AX foci in *Nex-Cre/Polβ^fl/fl^* mice (Figure 2). These results suggest the possibility that an abnormal level of SSBs in *Nex-Cre/Polβ^fl/fl^* mice leads to DSB formation.

**Fig. 3.**
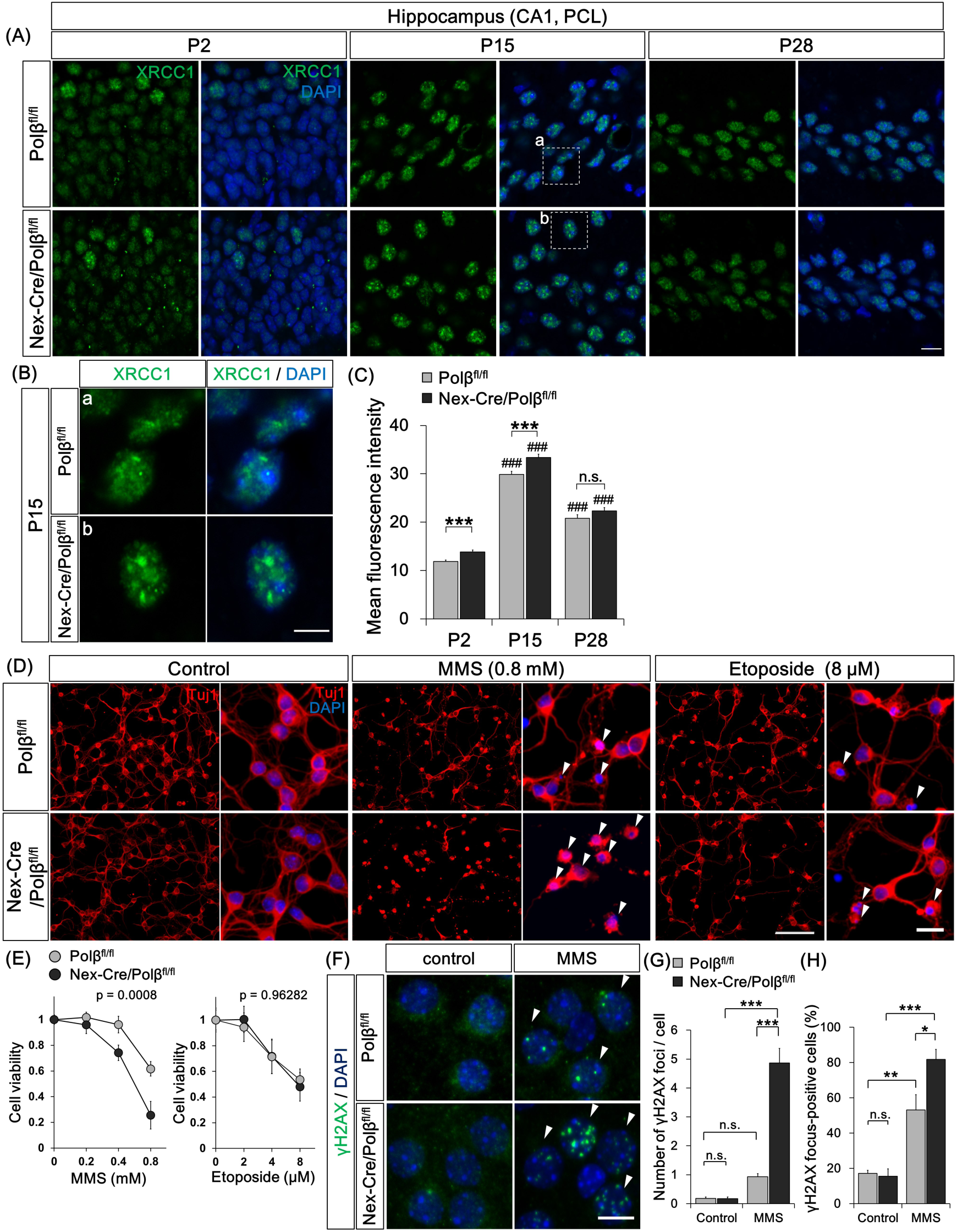
Polβ is required for SSB repair in postmitotic neurons. (A) Immunohistochemistry was performed with anti-XRCC1 antibody in CA1 pyramidal cell layers of *Nex-Cre/Polβ^fl/fl^* and *Polβ^fl/fl^* hippocampus at P2, P15, and P28. Scale bar, 10 µm. (B) Magnified images of the boxed areas in (A) are shown. Scale bar, 5 µm. (C) Histogram shows the mean XRCC1 fluorescence intensity in DAPI-stained nuclei of *Nex-Cre/Polβ^fl/fl^* (n = 576, 295, and 234 cells) and *Polβ^fl/fl^* (n = 643, 282, and 256 cells) hippocampal CA1 cells at P2, P15, and P28. Data are the mean ± SEM. Significant difference from *Polβ^fl/fl^*mice: ***p < 0.001, ANOVA with Tukey’s post-hoc test. Significant difference between age groups: ^###^p < 0.001, ANOVA with Tukey’s post-hoc test. (D) Primary cultured neurons from E16.5 *Nex-Cre/Polβ^fl/fl^* or *Polβ^fl/fl^* cortex were treated with MMS or etoposide for 1 h at 3–4 days *in vitro* (DIV), and fixed after 24 h recovery. Immunocytochemistry was then performed with anti-Tuj1 antibody. Arrowheads indicate dying pyknotic cells. Magnified images are shown in the right panels. Scale bars, 100 and 20 µm. (E) Quantitative analysis of cell viability of *Nex-Cre/Polβ^fl/fl^* and *Polβ^fl/fl^* cortical neurons treated with MMS or etoposide for 1 h. Data are mean ± SEM from three independent experiments. Significant difference: p-values (repeated measures ANOVA) are indicated. (F) Primary cultured neurons from E16.5 *Nex-Cre/Polβ^fl/fl^* or *Polβ^fl/fl^* cortex were treated with MMS for 1 h at 14 DIV and immunocytochemistry was performed with anti-γH2AX antibody. Arrowheads indicate γH2AX focus-positive cells. (G, H) Histograms show quantitative analysis of the mean number of γH2AX foci in each nucleus (G) and the percentage of γH2AX focus-positive cells (H). Data are mean ± SEM from control or MMS-treated *Nex-Cre/Polβ^fl/fl^* (n = 79 cells, n = 80 cells) and *Polβ^fl/fl^* (n = 71 cells, n = 86 cells) neurons in three independent experiments. Significant difference: *p < 0.05, **p < 0.01, and ***p < 0.001, ANOVA with Tukey’s post-hoc test.

The sensitivity of Polβ-deficient neurons to specific DNA-damaging agents was examined to reveal the role of Polβ in BER and DSBR. Primary cultured neurons from E16.5 control or *Nex-Cre/Polβ^fl/fl^* mouse cortex were treated with methylmethanesulfonate (MMS), which induces base damage (Beranek, 1990; Kulkarni et al, 2008; Sobol et al, 1996), or etoposide, an inhibitor of topoisomerase II that induces DSBs (Dobbin et al, 2013; Ross et al, 1984). Polβ-deficient neurons showed higher sensitivity to MMS than control (Figure 3D, E). In contrast, there was no significant difference following the etoposide treatment (Figure 3D, E). In addition, γH2AX focus formation after MMS treatment was significantly increased in neuronal cultures from *Nex-Cre/Polβ^fl/fl^* mice compared to those from *Polβ^fl/fl^* mice (Figure 3F-H). Taken together, these results demonstrate that Polβ is required for BER rather than DSBR, suggesting that highly accumulated SSBs are converted to DSBs in Polβ-deficient neurons during postnatal development.

### Loss of Polβ in active DNA demethylation causes DSBs in developing neurons

To examine the possibility that active DNA demethylation was a cause of the DSB formation in Polβ-deficient neurons during postnatal development (Lister et al, 2013; Sharma et al, 2016; Wu & Zhang, 2017), developmental changes in 5mC and 5hmC levels were quantified between P2 and P28. Immunoblot analysis with specific antibodies revealed that both 5mC and 5hmC levels decreased strongly in control *Polβ^fl/fl^* hippocampus between P2 and P15 (Figure 4A, B), during which the extent of SSB and DSB formation increased (Figure 2, 3A-C). This suggests that many DNA demethylation reactions occur on the genome during this period.

**Fig. 4.**
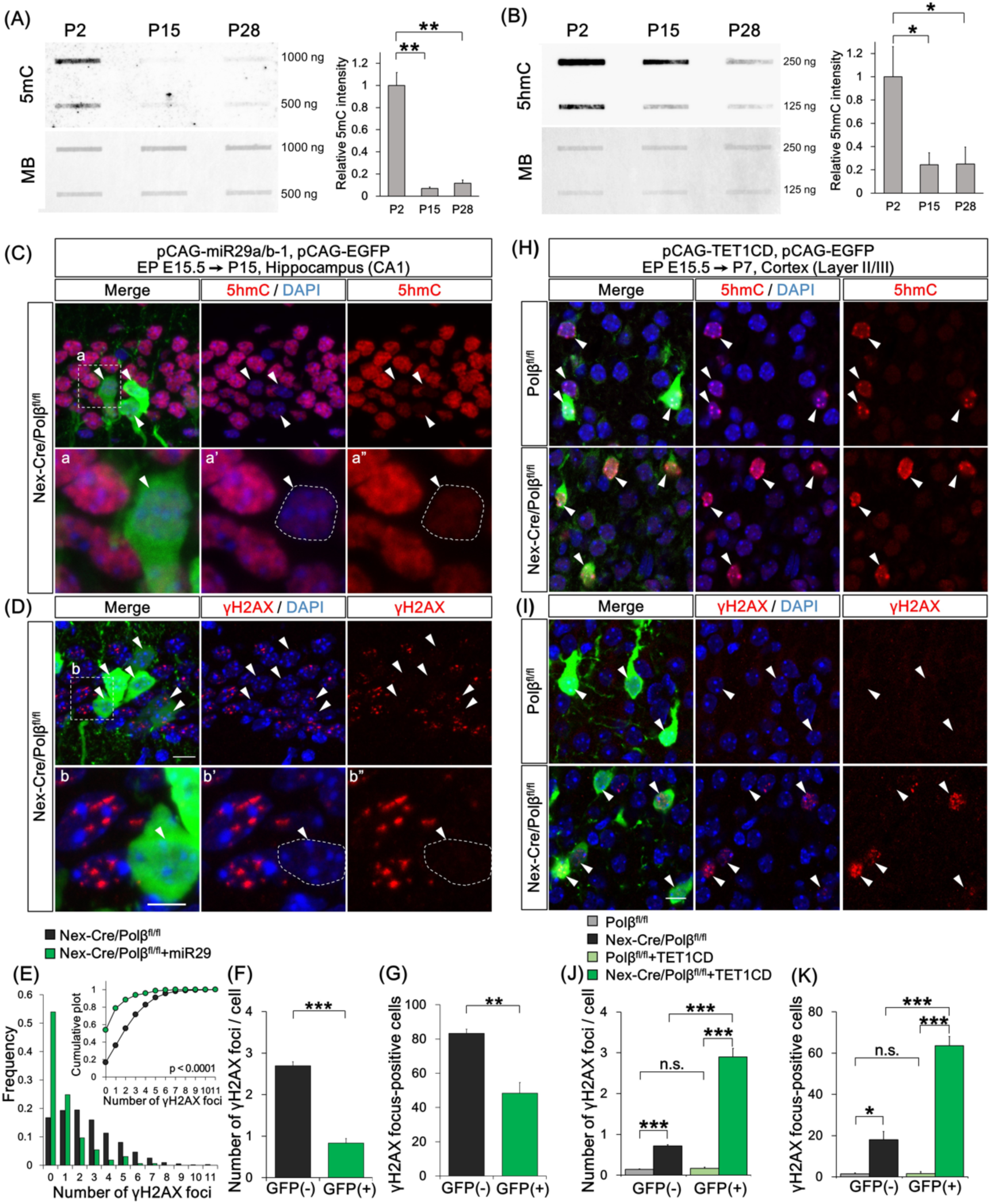
Loss of Polβ in active DNA demethylation causes DSBs in neurons. (A, B) Immunoblot analyses show amounts of 5mC (A) and 5hmC (B) in genomic DNA from P2, P15, and P28 *Polβ^fl/fl^* hippocampus. The membranes were also stained with methylene blue (MB) as a loading control. The relative intensity of 5mC and 5hmC was quantified. Data are mean ± SEM from three different brains. Significant difference: *p < 0.05 and **p < 0.01, ANOVA with Tukey’s post-hoc test. (C–G) Hippocampal CA1 neurons were cotransfected with pCAG-miR29a/b-1 and pCAG-EGFP by *in utero* electroporation at E15.5 and analyzed at P15. Immunohistochemistry was performed with anti-5hmC (C), -γH2AX (D), and -GFP antibodies in *Nex-Cre/Polβ^fl/fl^* hippocampus. (C, D) Magnified images of the dashed-line boxed areas in the upper image are shown in the the lower images. The dashed lines in the lower images mark the perimeter of the nucleus. Arrowheads indicate EGFP-positive transfected cells. Scale bars, 10 (the upper) and 5 (the lower) µm. (E) Distribution histogram shows the number of γH2AX foci in the transfected (GFP(+), n = 137 cells) and the surrounding untransfected (GFP(-), n = 522 cells) nuclei of *Nex-Cre/Polβ^fl/fl^* hippocampus. The Kolmogorov-Smirnov (KS) test shows the significant difference between GFP(-) and GFP(+) cells. (F, G) Histograms show the average number of γH2AX foci (F) and percentage of focus-positive cells (G) in the GFP(+) and the surrounding GFP(-) nuclei of *Nex-Cre/Polβ^fl/fl^* CA1 cells. Data are the mean ± SEM from three different brains. Significant difference: **p < 0.01 and ***p < 0.001, Student’s t-test. (H–K) Cortical upper layer neurons were cotransfected with pCAG-TET1CD and pCAG-EGFP by *in utero* electroporation at E15.5 and analyzed at P7. Immunohistochemistry was performed with anti-5hmC (H), -γH2AX (I) and -GFP antibodies in *Nex-Cre/Polβ^fl/fl^* and *Polβ^fl/fl^* cortex. Arrowheads indicate EGFP-positive transfected cells. Scale bar, 10 µm. (J, K) Histograms show the average number of γH2AX foci (J) or percentage of focus-positive cells (K) in the GFP(+) and surrounding GFP(-) nuclei of *Nex-Cre/Polβ^fl/fl^* (n = 160 cells, n = 950 cells) and *Polβ^fl/fl^* (n = 161 cells and n = 966 cells) cortex. Data are the mean ± SEM from three different brains. Significant difference: *p < 0.05, **p < 0.01, and ***p < 0.001, ANOVA with Tukey’s post-hoc test.

We examined whether inhibition of the active DNA demethylation process could affect DSB formation in Polβ-deficient hippocampal neurons *in vivo*. Overexpression of microRNAs miR29a and miR29b-1 has been reported to inhibit expression of several genes involved in DNA methylation and demethylation processes, resulting in a decrease in 5hmC level in transfected cells (Cheng et al, 2013; Hysolli et al, 2016). The miRNA expression vector was transfected into CA1 neurons in *Nex-Cre/Polβ^fl/fl^* hippocampus using an *in utero* electroporation technique. miR29a/b-1 efficiently decreased the 5hmC level in the transfected hippocampal neurons (Figure 4C). The numbers of γH2AX foci and of focus-positive cells in γH2AX foci were significantly lower in the transfected neurons than in the surrounding untransfected neurons of P15 *Nex-Cre/Polβ^fl/fl^* hippocampus (Figure 4D-G), indicating that inhibition of active DNA demethylation suppresses DSB formation.

Conversely, we tested whether induction of active demethylation promotes DSB formation in Polβ-deficient cortical neurons. TET1 catalytic domain (TET1CD), which induces 5hmC more efficiently than full-length TET1 (Tahiliani et al, 2009), was overexpressed in cortical neurons using *in utero* electroporation. As expected, the 5hmC level increased in both the transfected *Polβ^fl/fl^* and *Nex-Cre/Polβ^fl/fl^* cortical neurons at P7 (Figure 4H). The γH2AX foci were increased in the transfected neurons compared to the untransfected neurons of *Nex-Cre/Polβ^fl/fl^* cortex, but not of *Polβ^fl/fl^* cortex (Figure 4I). Quantitative analysis also demonstrated that both parameters of γH2AX foci were significantly increased in the transfected neurons relative to the surrounding neurons in *Nex-Cre/Polβ^fl/fl^*cortex, indicating that induction of active DNA demethylation is sufficient to promote DSB formation in Polβ-deficient cortical neurons (Figure 4J, K).

Finally, to modulate the endogenous active DNA demethylation process, cultured cortical neurons from E16.5 *Nex-Cre/Polβ^fl/fl^* and *Polβ^fl/fl^* mice were treated with vitamin C, which induces TET1 activity (Blaschke et al, 2013). Immunocytochemistry with anti-5hmC antibody showed an apparent increase in 5hmC level following 24-h vitamin C treatment in both control and Polβ-deficient neuronal nuclei (Figure S3A). Analysis of the DSB formation under this culture condition demonstrated that both the number of γH2AX foci and the proportion of focus-positive cells were significantly increased in *Nex-Cre/Polβ^fl/fl^* neurons but not in controls (Figure S3B-D). Together, these results suggest that active DNA demethylation is a primary cause of DSB formation in Polβ-deficient neurons during postnatal development.

### Polβ deficiency affects gene expression and dendrite morphology of hippocampal neurons during postnatal development

To investigate the possibility that active DNA demethylation defects and/or DSB formation alter gene expression in Polβ-deficient neurons, RNA-seq analysis was performed with RNA extracted from P15 control *Polβ^fl/fl^* and *Nex-Cre/Polβ^fl/fl^* hippocampus. Overall, 219 genes were found to be downregulated, and 199 upregulated, in the *Nex-Cre/Polβ^fl/fl^* hippocampus compared to the control (Figure 5A; n = 3, p < 0.05, fold change > 1.2). A functional annotation analysis of these 418 differentially expressed genes (DEGs) was performed using Ingenuity Pathway Analysis (IPA) software (Figure S4A-C). Genes related to nervous system development and function (p = 1.41×10^-2^) and to neurological diseases (p = 1.26 × 10^-4^), in addition to cancer, were significantly enriched in the DEGs (Figure 5B, C). In the canonical pathways identified by the IPA, signaling pathways related to cell cycle regulation, DNA damage response, and cancer cells were primarily suggested (Figure 5D), which may reflect a response to DSB formation in Polβ-deficient neurons. In addition, among the top hits, the assembly of RNA polymerase I complex (p = 0.012), the opioid signaling pathway (p = 0.013), and the Wnt/Ca+ pathway (p = 0.016), which are known to relate to neuronal development and learning and memory in the hippocampus, were included (Capitano et al, 2016; Inestrosa & Varela-Nallar, 2015; Williams et al, 2001). These results indicate that Polβ deficiency affects the regulation of genes involved in neuronal development and function. Furthermore, a marked similarity was identified in the gene expression profiles between the Polβ-deficient hippocampus and TET3 shRNA-transfected hippocampal neurons (Yu et al, 2015) using Illumina correlation engine software (Figure S4D), suggesting that some of the overlapped genes are under the control of active DNA demethylation.

**Fig. 5.**
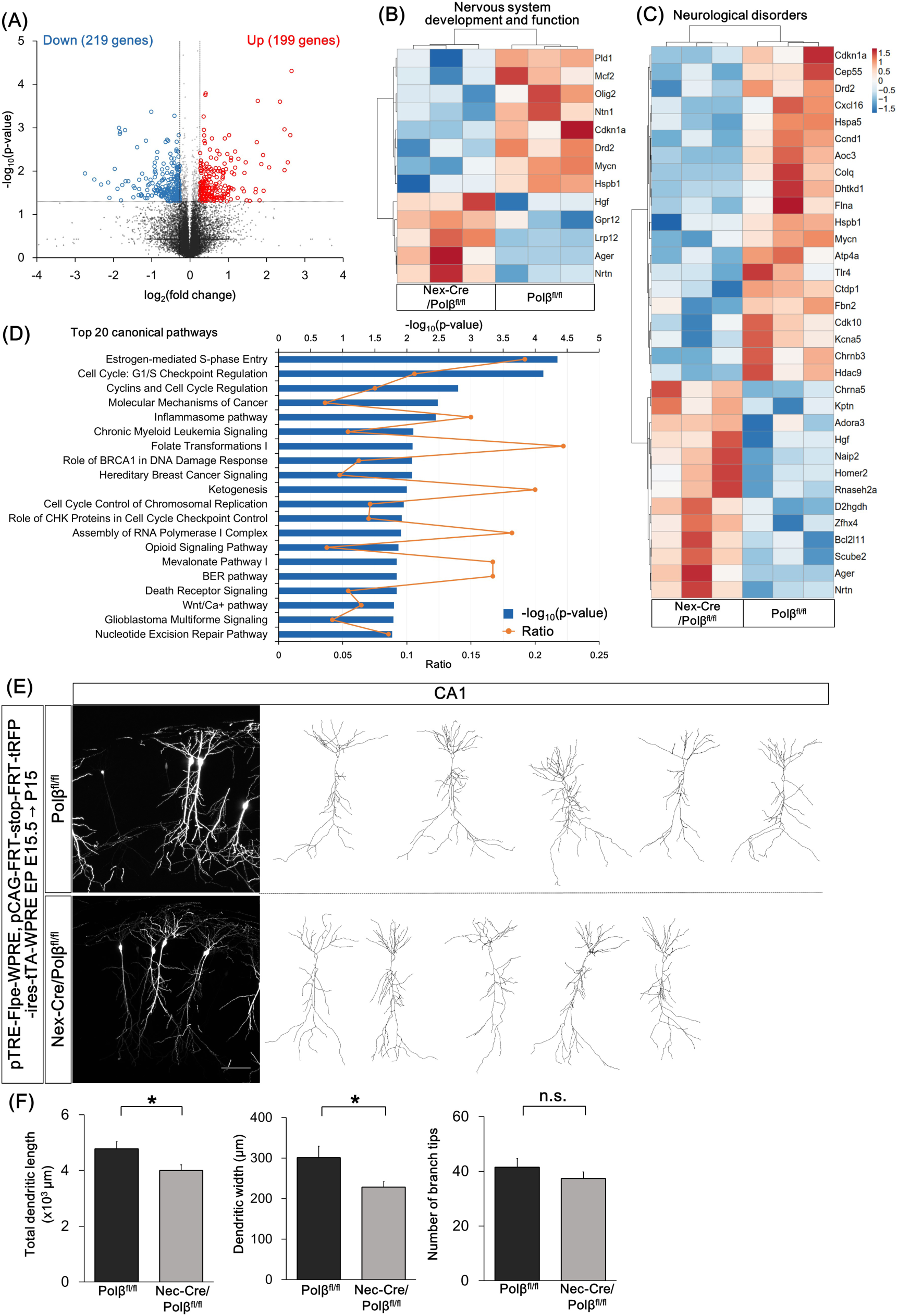
Polβ deficiency affects gene expression and dendrite morphology of hippocampal neurons during postnatal development. (A) Volcano plot of RNA-seq data from *Polβ^fl/fl^* and *Nex-Cre/Polβ^fl/fl^* P15 hippocampus (n = 3). Differentially expressed genes (DEGs, *Nex-Cre/Polβ^fl/fl^* vs *Polβ^fl/fl^*, p < 0.05, fold change > 1.2) are highlighted in blue (down) or red (up). (B, C) Hierarchical clustering with mean-centered log_2_-FPKM of DEGs related to nervous system development and function (B) or neurological disorders (C). Rows and columns represent genes and samples, respectively. (D) Top 20 canonical pathways predicted by Ingenuity Pathway Analysis. (E) Hippocampal CA1 neurons were cotransfected with FLPe-based Supernova vectors by *in utero* electroporation at E15.5 and analyzed at P15. Immunohistochemistry was performed with anti-tRFP antibody in *Nex-Cre/Polβ^fl/fl^* and *Polβ^fl/fl^* hippocampus. Examples of z-projected dendritic morphology of CA1 neurons traced with tRFP labeling are shown. Scale bar, 100 µm. (F) Quantitative analysis for total dendritic length, dendritic width, and number of branch tips. Data are mean ± SEM from *Polβ^fl/fl^* (n = 18 cells of 5 animals) and *Nex-Cre/Polβ^fl/fl^* (n = 20 cells of 4 animals) mice. Significant difference: *p < 0.05, ANOVA.

The altered gene expression in the *Nex-Cre/Polβ^fl/fl^* hippocampus may affect development of hippocampal neurons. Dendritic morphology of CA1 pyramidal neurons was examined in the *Nex-Cre/Polβ^fl/fl^* and *Polβ^fl/fl^* hippocampus. To visualize the morphology of individual neurons, sparse cell labeling was performed with the Supernova system using *in utero* electroporation (Luo et al, 2016; Mizuno et al, 2014). While both apical and basal dendrites of *Nex-Cre/Polβ^fl/fl^* CA1 neurons appeared to be similar to those in the control (Figure 5E), dendritic width (*Polβ^fl/fl^* vs *Nex-Cre/Polβ*^fl/fl^; 301 ± 28 µm, 228 ± 13 µm, p = 0.0204, ANOVA) and total dendrite length (*Polβ^fl/fl^* vs *Nex-Cre/Polβ*^fl/fl^; 4775 ± 263 µm, 3997 ± 207 µm, p = 0.0248, ANOVA) were significantly lower in *Nex-Cre/Polβ^fl/fl^* than in *Polβ^fl/fl^* neurons (Figure 5F). These results suggest that Polβ is required for dendrite formation in the developing hippocampus.

### *Nex-Cre/Polβ^fl/fl^* mice show impaired spatial reference memory and contextual fear memory

To further examine the involvement of Polβ in neuronal functions, *Nex-Cre/Polβ^fl/fl^* mice and their littermates were subjected to a comprehensive behavioral test battery (Shoji et al, 2018). Significant behavioral differences between control and mutant mice were found in several behavioral tests (Supplemental Table (Table S) 1). Notably, in the Barnes maze test, which is widely used for assessing spatial learning and memory, the number of errors to reach the target was significantly larger in *Nex-Cre/Polβ^fl/fl^* mice than in control *Polβ^fl/fl^* mice (Figure 6A; p = 0.0003). Consistent with this, the traveling distance and the latency were also significantly different between the two groups (Figure 6B, C). In the probe test at one day after final trial of the acquisition test, *Nex-Cre/Polβ^fl/fl^* mice spent significantly less time around their targets compared to the control (Figure 6D, 34.6 ± 3.9%, 20.7 ± 2.1%, p = 0.0031), confirming impaired spatial learning and memory due to distal environmental cues. This behavior was observed at one month after the acquisition test (Table S1). In the contextual and cued fear conditioning test, which is used to assess fear memory, no difference was found in freezing time during the conditioning phase and at one day after the conditioning (Figure 6E, Table S1). However, *Nex-Cre/Polβ^fl/fl^* mice showed a shorter freezing time in the contextual test at one month after the conditioning, but not in the cued test (Figure 6F, G). In addition, *Nex-Cre/Polβ^fl/fl^* mice also showed reduced anxiety behavior in the elevated plus maze (Figure 6H, I). Taking into consideration that spatial memory, contextual fear memory, and anxiety behavior are dependent on hippocampus function (Jimenez et al, 2018; Kim & Fanselow, 1992; Koopmans et al, 2003), these results suggest that the lack of Polβ tends to impair hippocampus-dependent functions.

**Fig. 6.**
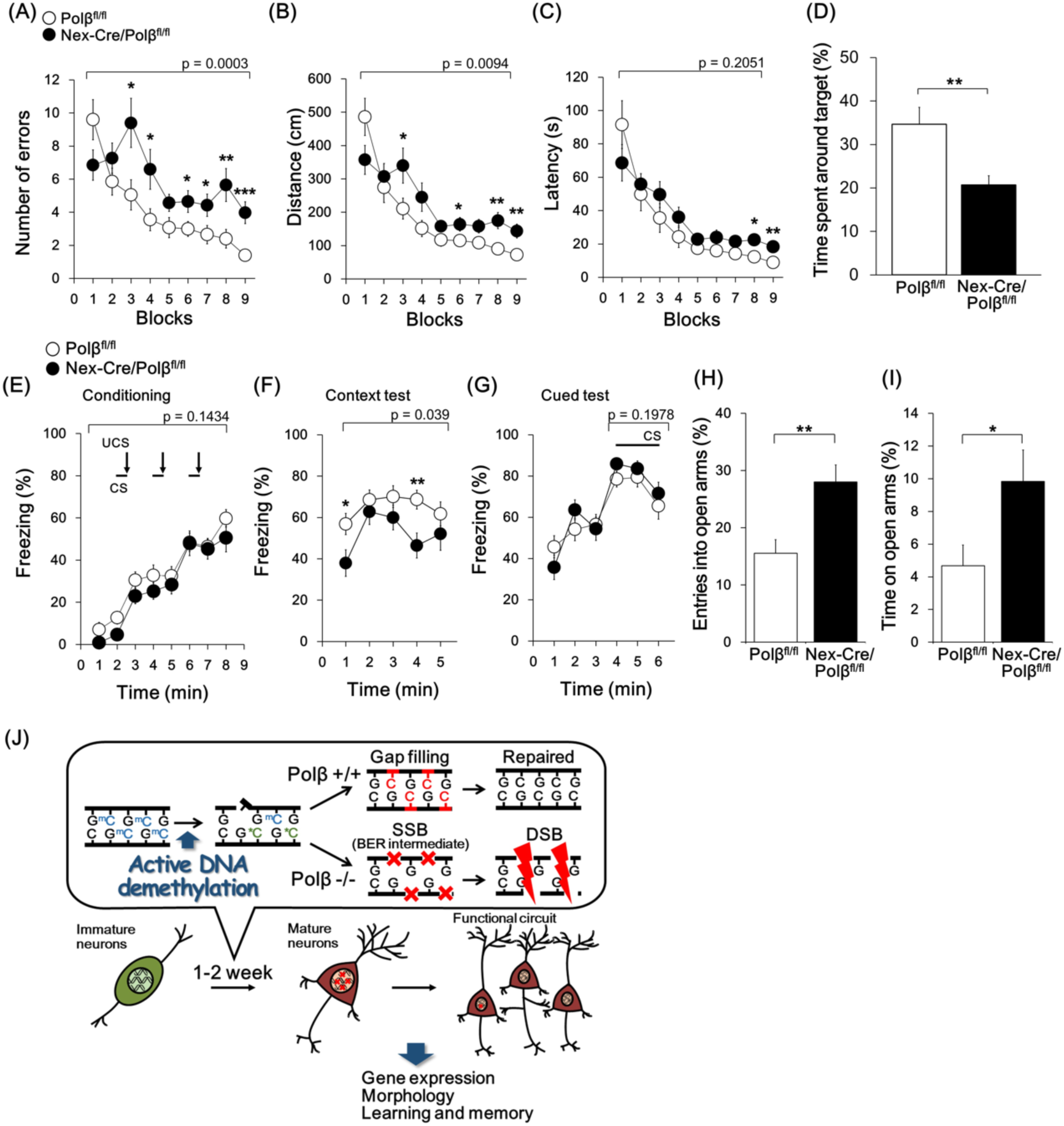
*Nex-Cre/Polβ^fl/fl^* mice show impaired spatial reference memory and contextual fear memory. (A–D) The Barnes maze test was performed for *Nex-Cre/Polβ^fl/fl^* (n = 20) and *Polβ^fl/fl^* (n = 20) mice. Quantitative analysis of the number of errors (A), distance traveled (B), and latency (C) before reaching the target hole. Data are mean ± SEM. Significant differences: p-values of repeated measures ANOVA are indicated. *p < 0.05, **p < 0.01, and ***p < 0.001, ANOVA for 2 blocks of trials. (D) Histogram shows percentage of time spent around the target in the probe test at one day after the acquisition test. Data is mean ± SEM. Significant differences: **p < 0.01, ANOVA. (E–G) Quantitative analysis of freezing behavior (%) in the conditioning session (E), contextual (F) and cued (G) test in the fear conditioning test. Data is mean ± SEM from *Nex-Cre/Polβ^fl/fl^* (n = 20) and *Polβ^fl/fl^* (n = 20) mice. Significant differences: p-values of repeated measures ANOVA are indicated. *p < 0.05, **p < 0.01, ANOVA for each duration. (H, I) Histograms show quantitative analysis of entries into open arms (H) and time spent in open arms (I) in the elevated plus maze test. Data are mean ± SEM from *Nex-Cre/Polβ^fl/fl^* (n = 20) and *Polβ^fl/fl^* (n = 20) mice. Significant difference, *p < 0.05, **p < 0.01, ANOVA. (J) Proposed model of Polβ-dependent active DNA demethylation during postnatal neuronal development.

## Discussion

The present study demonstrated that the loss of Polβ leads to DSB accumulation in developing hippocampal neurons, and to a lesser extent in cortical neurons, which is attributable to a failure of active DNA demethylation. The DSB accumulation in Polβ-deficient neurons did not induce apoptosis, but affected gene expression and dendritic morphology in hippocampal neurons. Furthermore, behavioral tests demonstrated that the loss of Polβ impaired hippocampal-dependent function. These results suggest that genome maintenance by Polβ contributes to hippocampal neuronal differentiation and functional circuit formation via epigenetic regulation of gene expression (Figure 6J).

### Polβ-dependent BER is involved in active DNA demethylation during postnatal development of the nervous system

Our results show that Polβ plays a role in an active DNA demethylation process in postmitotic neuronal development (Figure 4), giving a first insight into its function in epigenome regulation *in vivo* (Weber et al, 2016). In the case of DNA demethylation in postmitotic cells, the TET-dependent active process involving BER seems to be a major reaction because the passive process coupled with DNA replication is dysfunctional (Schuermann et al, 2016; Wu & Zhang, 2017). Indeed, the decrease in DNA methylation level was paralleled by accumulation of SSBs, which may be BER intermediates, in the developing hippocampal genome (Figure 3A, B, 4A, and B). The timing between P2 and P15 is roughly consistent with previous studies (Lister et al, 2013; Simmons et al, 2013). Data in the culture experiment (Figure S3A) suggest that vitamin C is also important to promote active DNA demethylation during this period (Blaschke et al, 2013).

Moreover, Polβ deficiency altered expression of neuronal genes that are involved in transcription regulation (Olig2, Hdac9), the cytoskeleton (Flna, Homer2), and synaptic transmission (Drd2, Chrna5) (Figure 5A-C). The altered gene expression patterns were observed consistently in all tested samples of hippocampus in *Nex-Cre/Polβ^fl/fl^* mice, meaning that the alterations are determinate rather than stochastic, as is different from spontaneous DNA base damage (Figure 5A-C). Expression profiles of TET3 shRNA knockdown hippocampal neurons are also similar to those of *Nex-Cre/Polβ^fl/fl^* hippocampus (Figure S4D) (Rudenko et al, 2013, Yu et al, 2015). These results support the notion that Polβ contributes to specific gene regulation via active DNA demethylation during hippocampal development. The greater abundance of DSBs in hippocampal than in cortical neurons may reflect distinct gene expression profiles during this period (Figure 1). Therefore, it is likely that Polβ-dependent active DNA demethylation is involved in epigenetic regulation of nervous system development, although we need more studies focusing on neuronal cell types, developmental stages, and neural activities (Gontier et al, 2018; Kaas et al, 2013; Li et al, 2014; Yu et al, 2015; Zhu et al, 2016).

### Genome instability arising from Polβ deficiency leads to long-lasting effects on the genome and epigenome, rather than to apoptosis

We found that Polβ is required for the genome stability of postmitotic neurons (Figure 1). Combining this observation with previous studies, Polβ-dependent BER appears to play a key role in suppressing DSB formation at two distinct developmental stages, namely neurogenesis and postnatal neuronal differentiation. Why does the loss of Polβ induce DSBs in developing neurons? Polβ deficiency generally increases nicks and/or gaps as SSBs in the genome (Sobol et al, 1996). Indeed, SSBs increased significantly in Polβ-deficient neurons *in vivo* (Figure 3A–C). Although DNA replication can promote DSB formation from a single SSB in the case of mitotic neural progenitors (Kuzminov, 2001; Onishi et al, 2017), accumulation of SSBs may directly induce DSBs in postmitotic neurons. Supporting this notion, base damage introduced by MMS treatment induced DSBs in Polβ-deficient neuronal cultures (Figure 3F–H). On the other hand, neuronal apoptosis was undetectable in *Nex-Cre/Polβ^fl/fl^* hippocampal neurons (Figure S2C). Induction of apoptosis seems to be dependent on not only the extent of DSBs but also p53 pathway activity (Chong et al, 2000; Sugo et al, 2004).

The collapse of the active DNA demethylation process due to loss of Polβ causes DSB formation in postmitotic neurons (Figure 1). However, the extent of γH2AX foci in *Nex-Cre/Polβ^fl/fl^* mice declined after a peak at P15 (Figure 2). This suggests that DSBs in Polβ-deficient neurons are repaired by the DSBR pathway. Considering that postmitotic neurons only have NHEJ activity, which is error-prone DNA repair compared to homologous recombination in mitotic cells, mutations such as insertions/deletions are likely introduced into the DSB sites (Lieber, 2010). It will be interesting to determine whether DSBs and/or DSB-induced *de novo* somatic mutations are intensively introduced into mC-rich enhancer and promoter regions in neuronal genes. Indeed, the human neuronal genome and epigenome are drastically altered in the developing brain and this probably has a long-lasting influence on the brain into adulthood. (Cai et al, 2014; Lodato et al, 2015; Rehen et al, 2005; Stroud et al, 2017; Wei et al, 2016). Recent work has revealed that such alterations is increased in psychiatric diseases, although the underlying mechanism remains uncertain (Bundo et al, 2014; Fromer et al, 2014; Iossifov et al, 2014; McConnell et al, 2013). Our findings indicate that active DNA demethylation-associated DNA damage is a potential cause of *de novo* somatic mutations and an aberrant epigenome in brain developmental disorders.

### The role of Polβ in structure and function of the cerebral cortex

Polβ plays a role in the molecular bases underlying dendrite formation (Figure 5E, F). To date, TET1 and TET3 have also been shown to be involved in synaptic excitability and plasticity (Rudenko et al, 2013, Yu et al, 2015). These data suggest that DNA demethylation is necessary for neuronal development. On the other hand, it has also been demonstrated that the DNA methyltransferases Dnmt1, Dnmt3a, Dnmt3b, and the methyl-CpG binding protein MeCP2 are involved in dendritic arborization (Cohen et al, 2011; Feng et al, 2010; Golshani et al, 2005; Moretti et al, 2006; Zhou et al, 2006). Therefore, gene expression mediated by the bidirectional regulation of DNA methylation and demethylation may be crucial for neuronal development and circuit formation.

A comprehensive behavioral test with *Nex-Cre/Polβ^fl/fl^* mice showed a remarkable impairment in spatial reference memory and contextual fear memory (Figure 6A-G). This concurs with recent reports suggesting that active DNA demethylation is involved in learning and memory in adult mice (Gontier et al, 2018; Kaas et al, 2013; Li et al, 2014; Rudenko et al, 2013). However, considering that DSB accumulation in Polβ-deficient neurons is most prominent in early postnatal stages, we propose that DSBs and/or DSB-induced *de novo* mutations arising from the impairment of active DNA demethylation alter gene expression leading to circuit formation, and have a long-lasting influence on learning and memory. Indeed, the functional impairment was striking in the hippocampus (Figure 6, Table S1), in which DSBs accumulated extensively during the early postnatal stages (Figure 1, 2).

While active DNA demethylation is revealed as a potential source of DNA damage during early postnatal stages in *Nex-Cre/Polβ^fl/fl^* mice, we cannot completely rule out the possibility that oxidative stress causes DNA damage, leading to cognitive dysfunction in the adult (Wilson & McNeill, 2007). Indeed, Polβ-dependent impairment of cognitive function is reportedly accelerated in aged brains and Alzheimer’s disease models (Sykora et al, 2015). Moreover, the expression of DNA repair enzymes including Polβ gradually decreases with increased oxidative DNA damage associated with aging (Lu et al, 2004; Nowak et al, 1990; Rao et al, 2001; Wilson & McNeill, 2007). Therefore, although pathological changes with aging can affects the role of Polβ in neurons, our findings provide a key insight into its role during early postnatal development, which has long-lasting cognitive and behavioral outcomes.

## Methods

### Animals

All experiments were conducted under the guidelines for laboratory animals of the Graduate School of Frontier Biosciences, Osaka University. The protocol was approved by the Animal Care and Use Committee of the Graduate School of Frontier Biosciences, Osaka University and Fujita Health University. *Nex^Cre/+^Polβ^fl/fl^* (*Nex-Cre/Polβ^fl/fl^*) mice were generated as described previously (Goebbels et al, 2006; Gu et al, 1994; Iwasato et al, 2000; Onishi et al, 2017). Both male and female mice were used in all experiments expect RNA-seq analysis and the behavioral test. Noon of the day on which the vaginal plug was detected was designated as embryonic day 0.5 (E0.5) and the day of birth was designated as postnatal day 0 (P0). Genotyping was performed using the following primers: Polβ locus: 5’-CCACACCGAAGTCCTCTGAT-3’, 5’-AGGCTGGCCTCAGACTCATA-3’ and 5’-CTGGCTCACGTTCTTCTC-3’; Cre locus: 5’-GCAGAACCTGAAGATGTTCGCGAT-3’ and 5’-AGGTATCTCTGACCAGAGTCATCC-3’.

### Cell cultures

Pregnant mice were deeply anesthetized with pentobarbital (50 mg/kg, i.p.). Cortices were dissected from E16.5 embryos in ice-cold HBSS and then minced with fine scissors in PBS. The minced tissues were incubated with 0.125% trypsin and 0.02% EDTA in PBS for 5 min at 37°C, and then triturated thoroughly using a fire-polished Pasteur pipette. After centrifugation, the cells were resuspended in DMEM/F12 medium (Life Technologies) supplemented with B27 (Life Technologies) and 5% fetal bovine serum (Hyclone). A suspension containing 2.0×10^5^ cells was plated with culture medium on a 12 mm micro coverglass (Matsunami) in a multi-well dish (Thermo Scientific) coated with 0.1 mg/ml poly-L-ornithine (Sigma, P3655). The cultures were maintained at 37°C in an environment of 5% CO_2_ and humidified 95% air.

### Plasmids

pFN21AE2295, containing HaloTag-human TET1 cDNA, was purchased from Promega. To generate TET1 catalytic domain (TET1CD) expression vector pCAGGS-TET1CD, TET1CD was amplified from pFN21AE2295 by PCR with the following primers: TET1CD-F (+start) 5’-ATGGAACTGCCCACCTGCAGCTGTCT-3’ and TET1CD-R (+stop) 5’-TCAGACCCAATGGTTATAGGGCCCCG-3’. The PCR product was subcloned into pGEM-T Easy vector (A1360, Promega). An EcoRI-digested fragment containing TET1CD was then ligated to EcoRI-digested pCAGGS vector (Niwa et al, 1991). To generate pCAGGS-miR29a/b-1, the miR29a/b-1 cluster locus was amplified from mouse genomic DNA by PCR with the following primers: miR29a/b-1 F: 5’-TGTGTTGCTTTGCCTTTGAGAGGA-3’ and miR29a/b-1 R: 5’-CACATAGGGATAGTCACCTAGCCTG-3’; the product was subcloned into pGEM-T Easy vector. An EcoRI-digested fragment containing miR29a/b-1 locus was then ligated to EcoRI-digested pCAGGS vector. Supernova vectors pTRE-Flpe-WPRE (pK036) and pCAG-FRT-stop-FRT-tRFP-ires-tTA-WPRE (pK037) were kindly gifted from Dr. Iwasato (Luo et al, 2016). These vectors were cotransfected with pCAGGS-EGFP. All plasmids were purified with the PureLink HiPure Plasmid Maxiprep Kit (Invitrogen), and then dissolved in PBS.

### *In utero* electroporation

*In utero* electroporation was performed on E15.5 pregnant mice as previously described (Fukuchi-Shimogori & Grove, 2001, Tabata & Nakajima, 2001, Tomita, Kubo et al, 2011). Pregnant mice were deeply anesthetized with isoflurane (Wako Chemicals) using inhalation anesthesia equipment (KN-1071-1, Natsume). Plasmids (1-3 µg) were injected to the lateral ventricle with a glass micropipette connected to an injector (IM-30, Narishige). Electric pulses were delivered with disc-type electrodes (LF650P3 or LF650P5, BEX) connected to an electroporator (CUY21, BEX). Five 35-V pulses of 50 ms duration were applied at intervals of 950 ms.

### Pharmacological treatment

For a cell survival assay, cells at 3–4 DIV were incubated with culture medium containg 0–0.8 mM methylmethanesulfonate (MMS, Sigma, 129925) or 0–8 µM etoposide (Sigma, E1383) for 1 h, washed once with DMEM/F12 medium, and allowed to recover in conditioned medium for 24 h. To induce DNA base damage, cells at 14 DIV were treated with culture medium containing 0.4 mM MMS for 1 h, and then fixed. To induce Tet-dependent DNA demethylation, cells at 14 DIV were treated with culture medium containing 100 µg/ml L-ascorbic acid 2-phosphate (vitamin C, Sigma, 49752) for 24 h, and then fixed.

### Immunostaining

Mice were deeply anesthetized and perfused with phosphate buffered saline (PBS, pH 7.4) followed by 2% paraformaldehyde (PFA) in 0.1 M phosphate buffer (PB, pH 7.4). Their brains were postfixed in the same fixative on ice for 2 h, equilibrated with 25% sucrose in PBS, frozen in OCT compound (Sakura Finetech), and then sectioned at 10 or 20 µm using a cryostat (CM1850, Leica). The sections were permeabilized and blocked for 1 h at room temperature in buffer G (0.1 or 1.0% Triton X-100, 5% normal goat serum (Vector Laboratories) in PBS). They were then incubated at 4°C overnight with the following primary antibodies diluted in buffer G: rabbit polyclonal anti-cleaved caspase-3 (Asp175) (Cell Signaling, #9661) at 1:250, rabbit polyclonal anti-histone H2AX phosphor Ser139 (Active Motif, AR-0149-10) at 1:200, rabbit polyclonal anti-53BP1 (Gene Tex, GTX102595) at 1:200, rabbit monoclonal anti-XRCC1 (Abcam, ab134056) at 1:200, rat polyclonal anti-Ctip2 (Abcam, ab18465) at 1:800, rat monoclonal anti-GFP (Nacalai Tesque, GF090R) at 1:1000, and rabbit polyclonal anti 5-hydroxymethylcytosine (5hmC) (Active Motif, 39769) at 1:20000. Immunostaining of Polβ was performed as described previously (Onishi et al, 2017). For XRCC1 and 5hmC immunostaining, the sections were treated with 10 mM sodium citrate buffer (pH 6.0) for 10 min at 98°C using an autoclave. For co-immunostaining with anti-5hmC and -GFP antibodies, anti-GFP antibody was preincubated overnight at 4°C before the antigen retrieval step. Primary antibodies were detected by incubation with the secondary antibodies Alexa488-conjugated anti-rabbit IgG (A-11034, Invitrogen), Alexa488-conjugated anti-rat IgG (A-11006, Invitrogen), Alexa594-conjugated anti-rabbit IgG (A21207, Invitrogen), Cy3-conjugated anti-mouse IgG (AP192C, Millipore), Cy3-conjugated anti-rabbit IgG (AP182C, Millipore), and Cy3-conjugated anti-rat IgG (AP136C, Millipore), in all cases diluted at 1:400 in buffer G for 2 h at room temperature. Finally, the sections were mounted with a medium containing 0.1% 4’, 6-diamidine-2’-phenylindole (DAPI, Sigma), 1, 4-diazabicyclo [2, 2, 2] octane (Sigma) and 50 or 80% glycerol in 50 mM Tris-HCl (pH 8.0).

For morphological analysis of dendrites, mice were perfused with 4% PFA in 0.1 M PB (pH 7.4). The brains were postfixed in the same fixative for 24 h at 4°C, and equilibrated with 25% sucrose-PBS overnight at 4°C. The brains were cut into 200-µm coronal sections using a vibratome (DTK-1000, D.S.K.). The free-floating sections were permeabilized and blocked in buffer G for 1 h at room temperature. They were then incubated with primary antibodies, rat monoclonal anti-GFP (Nacalai Tesque, GF090R) at 1:2000 and rabbit polyclonal anti-tRFP (Evrogen, AB233) at 1:2000, in buffer G overnight at 4°C. The sections were washed three times with 0.1% Triton X-100 in PBS (PBST) for 1 h, and incubated with secondary antibodies in buffer G overnight at 4°C. The sections were then washed three times with PBST for 1 h and mounted with DAPI-containing mounting medium.

Cultured cells were fixed with 4% PFA in PBS for 10 min at room temperature, washed three times with PBS for 10 min, permeabilized, and blocked in buffer G. The cells were then incubated overnight at 4°C in buffer G with the following antibodies: mouse monoclonal anti-Tuj1 (R&D Systems, MAB1195) at 1:1000, rabbit polyclonal anti-histone H2AX phosphor Ser139 (Active Motif, AR-0149-10) at 1:200, and rabbit polyclonal anti 5-hydroxymethylcytosine (Active Motif, 39769) at 1:20000. For 5hmC immunostaining, the cells were incubated with 1 M HCl for 30 min at 37°C and washed three times with PBS for 30 min before permeabilization. Primary antibodies were detected by incubation for 2 h at room temperature in buffer G with the following secondary antibodies: Alexa488-conjugated anti rabbit IgG (A-11034, Invitrogen) at 1:400 and Cy3-conjugated anti-mouse IgG (AP192C, Millipore) at 1:400. The cells were mounted with DAPI-containing mounting medium.

### Image analysis

Fluorescence images were obtained by confocal microscopy (ECLIPSE FN with EZ-C1; Nikon) with 10×/0.3, 20×/0.75, and 40×/0.95 objective lenses (Nikon). All images were imported into ImageJ to adjust brightness and contrast. To acquire images of γH2AX and XRCC1 in DAPI-stained nuclei, confocal z-stack images were collected at 0.5-µm intervals with a 40× objective lens. For focus counting, noise in the images was removed by Gaussian filter and subtraction from the background, and foci were detected with the “Find Maxima” tool in ImageJ. To obtain images of RFP-labeled apical and basal dendrites in dorsal hippocampal CA1 regions, confocal z-stack images were collected at 1-µm intervals through the 200-µm sections using a 20× objective lens. Quantitative analysis of dendrite morphology was performed using the ImageJ plug-ins Simple Neurite Tracer and L-measure (Scorcioni et al, 2008).

### Immunoblot analysis

Genomic DNA was extracted using a DNeasy Blood & Tissue kit (QIAGEN). The DNA was eluted with TE buffer and stored at –30°C until required. The genomic DNA was denatured in 20 mM Tris-HCl (pH 8.0) for 10 min at 98°C and chilled on ice. Serially diluted DNA samples (1000, 500, 250, 125 ng/200 µl) were blotted onto a positively charged nylon membrane (Millipore, INYC00010) using a Bio-Dot slot blot apparatus (Bio-Rad). The membrane was air-dried, and UV-crosslinked using a CL-1000 Ultraviolet Crosslinker (UVP). The membrane was stained with 0.02% methylene blue (Nacalai Tesque) for 30 min at room temperature as a loading control. The membrane was then washed with Tris-buffered saline containing 0.1% Tween-20 (TBS-T) and blocked in 5% nonfat dry milk (Cell Signaling Technology, #9999) diluted with TBS-T for 1 h at room temperature. The membrane was incubated with the following primary antibodies diluted in 5% nonfat dry milk/TBS-T overnight at 4°C: rabbit polyclonal anti-5-hydroxymethylcytosine antibody (Active Motif, 39769) at 1:5000 and mouse monoclonal anti-5-methylcytosine antibody (Active Motif, 39649) at 1:2000. Primary antibodies were detected by incubation with the following secondary antibodies diluted in 5% skim milk in TBS-T for 2 h at room temperature: peroxidase-conjugated anti-rabbit IgG antibody (Jackson ImmunoResearch, 711-035-152) at 1:5000 and peroxidase-conjugated anti-mouse IgG (Nacalai Tesque, 01803-44) at 1:5000. The signal was visualized by chemiluminescence with ECL Select western blotting detection reagent (GE Healthcare) and imaged by LAS-3000UV mini (Fujifilm).

### RNA seq analysis

Total RNA was extracted from P15 *Polβ^fl/fl^* and *Nex-Cre/Polβ^fl/fl^* hippocampus using an RNeasy Plus Mini Kit (QIAGEN) following the manufacturer’s procedure. Library preparation was performed using a TruSeq stranded mRNA sample prep kit (Illumina) according to the manufacturer’s instructions. Whole-transcriptome sequencing was applied to the RNA samples with an Illumina HiSeq 2500 platform in a 75-base single-end mode. Illumina Casava ver.1.8.2 software was used for base calling. Sequenced reads were mapped to the mouse reference genome sequences (mm10) using TopHat ver. 2.0.13 in combination with Bowtie2 ver. 2.2.3 and SAMtools ver. 0.1.19. The number of fragments per kilobase of exon per million mapped fragments (FPKMs) was calculated using Cufflinks ver. 2.2.1. Differentially expressed genes were defined by fold change > 1.2, p < 0.05 (n = 3). Functional annotation and pathway analysis were performed with Ingenuity pathway analysis (QIAGEN). Correlation analysis of expression profiles was performed with the Illumina correlation engine software (Illumina).

### Behavioral test

Behavioral tests were carried out at Institute for Comprehensive Medical Science, Fujita Health University (Joint Usage / Research Center for Genes, Brain and Behavior accredited by MEXT). The comprehensive behavioral test was performed as described previously (Shoji et al, 2018) with adult (> 3 month) *Nex-Cre/Polβ^fl/fl^* mice and their littermate controls. In brief, the Barnes maze test was performed with a white circular surface, 1.0 m in diameter, with 12 holes equally spaced around the perimeter and elevated 75 cm from the floor. A black Plexiglas escape box was located under one of the holes and represented the target. The location of the target was consistent for each mouse but randomized between mice. The visual cues were in the four corners of the experimental room. One or two trials per day were performed. The number of errors, latency to reach the target, and distance traveled before mice first reached their target were automatically calculated by image analysis. One day or one month after the last training session, a probe test was performed without the escape box for 3 min, and time spent around each hole was measured and the ratio of time spent around the target / all holes was quantified.

In conditioning session of contextual and cued fear-conditioning test, each mouse was placed in a transparent acrylic chamber with a stainless steel grid floor (O’Hara & Co.) and allowed to explore for 2 min. White noise (55 dB) was then presented for 30 sec as a conditioned stimulus (CS). A mild footshock (0.3 mA, 2 sec) was presented as an unconditioned stimulus (UCS) during the last 2 sec of the CS. Three CS-UCS pairings were presented with a 2-min interval. One day or one month after the conditioning session, contest tests were conducted in the same chamber as conditioning. Cued tests with altered context were then conducted in a triangular and white opaque chamber which was located in a different room. In each test, freezing percentage and distance traveled in 1 min were quantified.

In the elevated plus maze, each mouse was placed in the central square of the maze, which consisted of two open arms (25 x 5 cm) and two closed arms (25 x 5 cm, with 15-cm-high transparent walls), and allowed to explore for 10 min. The number of total entries into the arms, percentage of entries into the open arms, and percentage of time spent in the open arms were quantified.

### Statistical analysis

In statistical analysis, the number of samples analyzed is given for each experiment. Significant differences were determined with Student’s t-test, one-way ANOVA with Tukey’s post-hoc test for multiple comparisons, or repeated-measures ANOVA. All statistical values are presented as mean value ± SEM. All data were analyzed using Excel 2013 (Microsoft), StatView 5.0.1 software (SAS Institute), and JMP (SAS Institute).

## Acknowledgments

We are grateful to Drs. K.A. Nave for the Nex-Cre mice, and to K. Rajewsky for Polβ flox mice. This work was supported by MEXT KAKENHI on Dynamic regulation of brain function by Scrap & Build system (No. 16H06460) to N.Y., by JSPS KAKENHI Grant Nos. 15K14350 and 17K07109 to N.S., 16H06276 (AdAMS) to N.S. and T.M., and by AMED-CREST to T.Y.

## Author contributions

T.M., T.Y., N.Y. and N.S. conceived and designed the research project. A.U., K.O., T.H., S.H. and N.S. performed the experiments and analyzed the data. A.U., T.M., T.Y., N.Y. and N.S. wrote the manuscript, which was discussed and critically edited by all coauthors.

## Figures and Figure Legends

**Supplemental Fig. 1.**
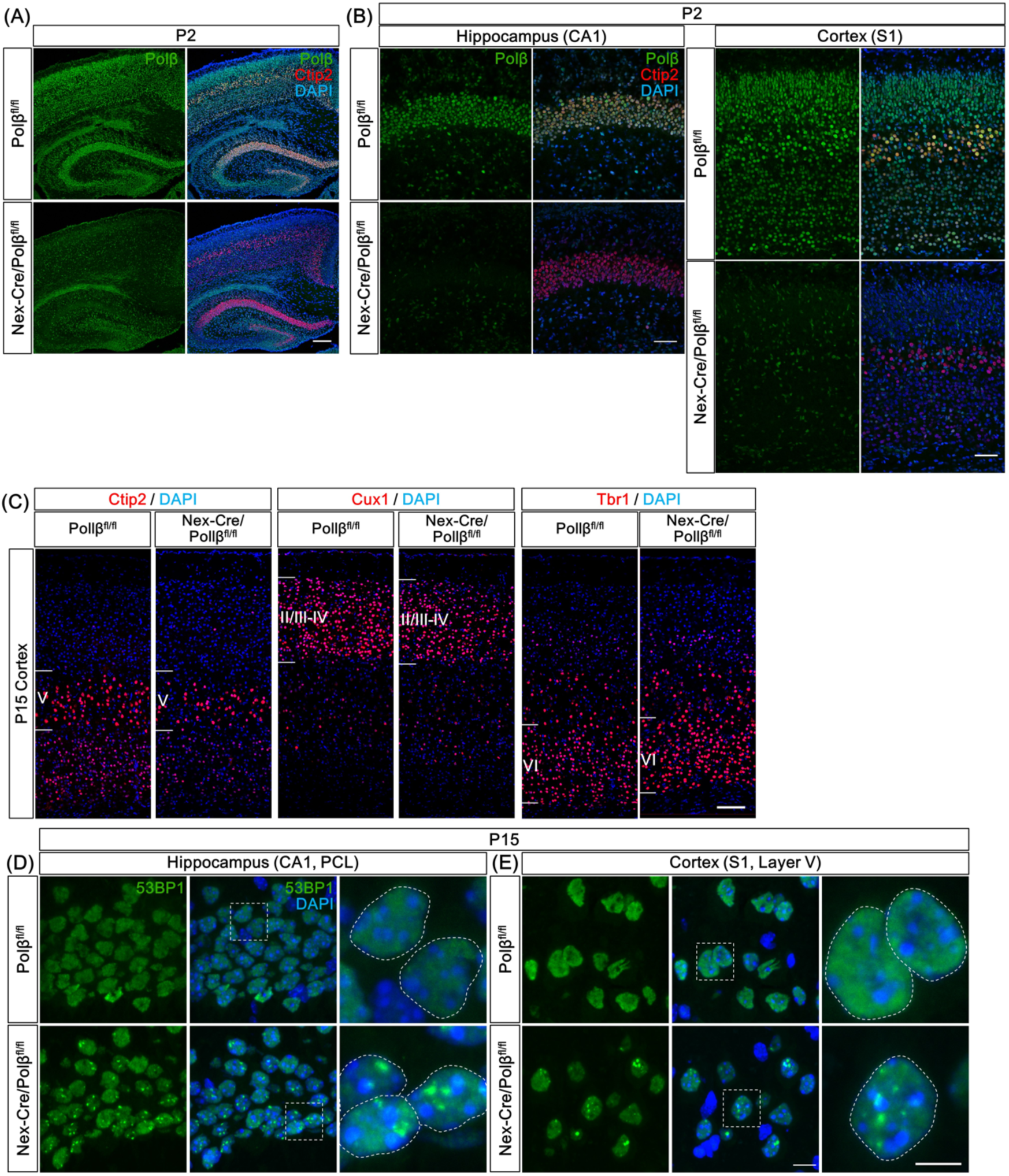
*Nex-Cre/Polβ^fl/fl^* mice exhibit DSB formation in postnatal hippocampus and cortex. (A) Immunohistochemistry was performed with anti-Polβ and -Ctip2 antibodies in P2 *Nex-Cre/Polβ^fl/fl^* and control *Polβ^fl/fl^* cortex and hippocampus. Scale bar, 400 µm. (B) Magnified images are hippocampal CA1 area and somatosensory area 1 (S1) in the cortex. Scale bar, 100 µm. (C) Immunohistochemistry was performed with anti-Ctip2, -Cux1, and -Tbr1 antibodies in P15 *Nex-Cre/Polβ^fl/fl^* and *Polβ^fl/fl^* cortex. Scale bar, 200 µm. (D) Immunohistochemistry was performed with anti-53BP1 antibody in P15 *Nex-Cre/Polβ^fl/fl^* and *Polβ^fl/fl^* cortex and hippocampus. Magnified images of the boxed areas are shown in the right panels. The dashed lines in the rightmost images mark the perimeter of the nucleus. Scale bars, 10 (the center) and 5 (the right) µm.

**Supplemental Fig. 2.**
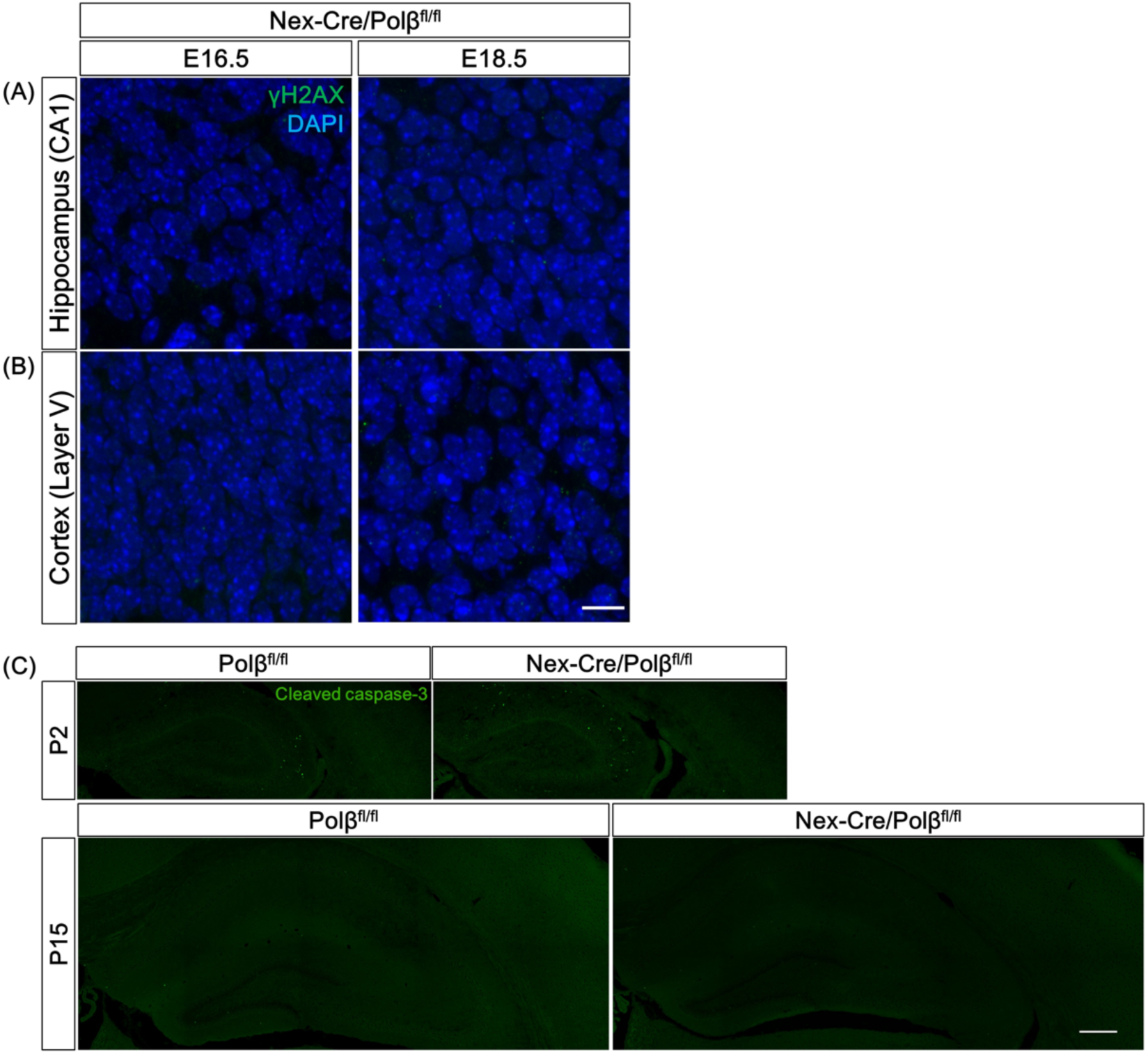
*Nex-Cre/Polβ^fl/fl^* mice exhibit the extent of apoptotic cells similar to that control hippocampus and cortex. (A, B) Immunohistochemistry was performed with anti-γH2AX antibody in *Nex-Cre/Polβ^fl/fl^* and *Polβ^fl/fl^* hippocampus (A) and cortex (B) at E16.5 and E18.5. Scale bar, 10 µm. (C) Immunohistochemistry was performed with anti-cleaved caspase 3 antibody in *Nex-Cre/Polβ^fl/fl^* and *Polβ^fl/fl^* hippocampus and cortex at P2 and P15. Scale bar, 400 µm.

**Supplemental Fig. 3.**
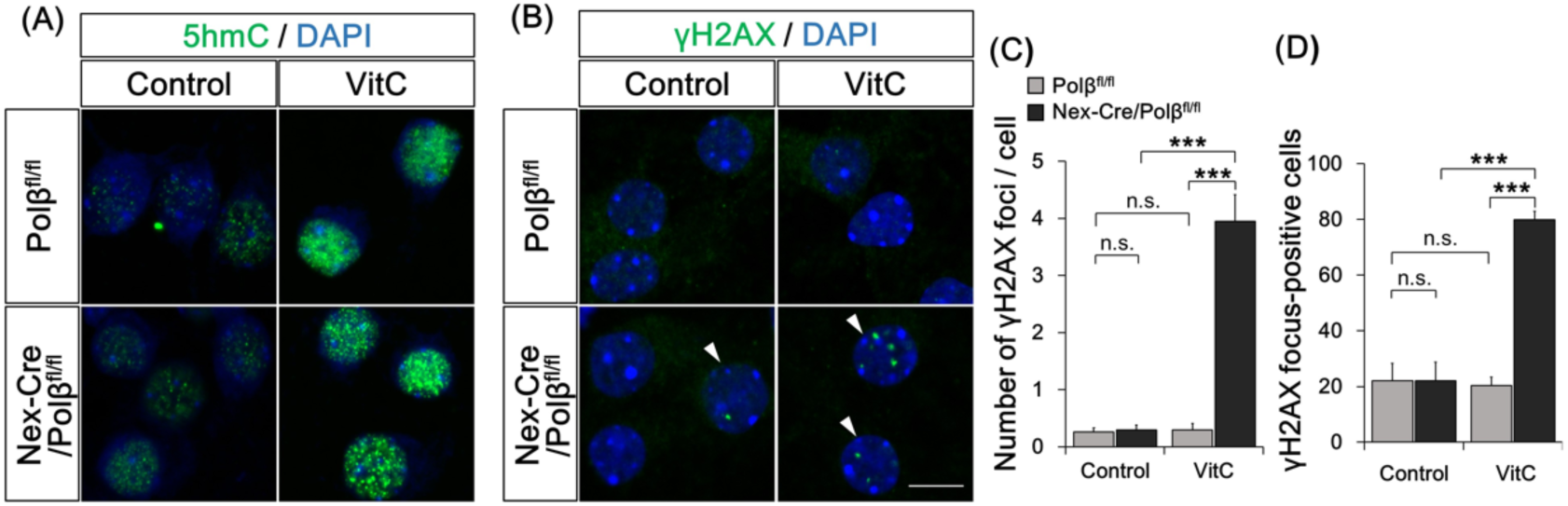
Loss of Polβ in active DNA demethylation causes DSBs in neurons. (A, B) Primary cultured neurons from E16.5 *Nex-Cre/Polβ^fl/fl^* or *Polβ^fl/fl^* cortex were treated with vitamin C (VitC) for 24 h at 14 DIV and immunocytochemistry was performed with anti-5hmC (A) or anti-γH2AX (B) antibodies. Arrowheads indicate γH2AX focus-positive cells. (C, D) Histograms show the mean number of γH2AX foci in each nucleus (C) and the percentage of γH2AX focus-positive cells. Data are the mean ± SEM from control or VitC-treated *Nex-Cre/Polβ^fl/fl^* (n = 48 cells, n = 62 cells) and *Polβ^fl/fl^* (n = 54 cells, n = 49 cells) cortical neurons in three independent experiments. Significant difference: ***p < 0.001, ANOVA with Tukey’s post-hoc test.

**Supplemental Fig. 4.**
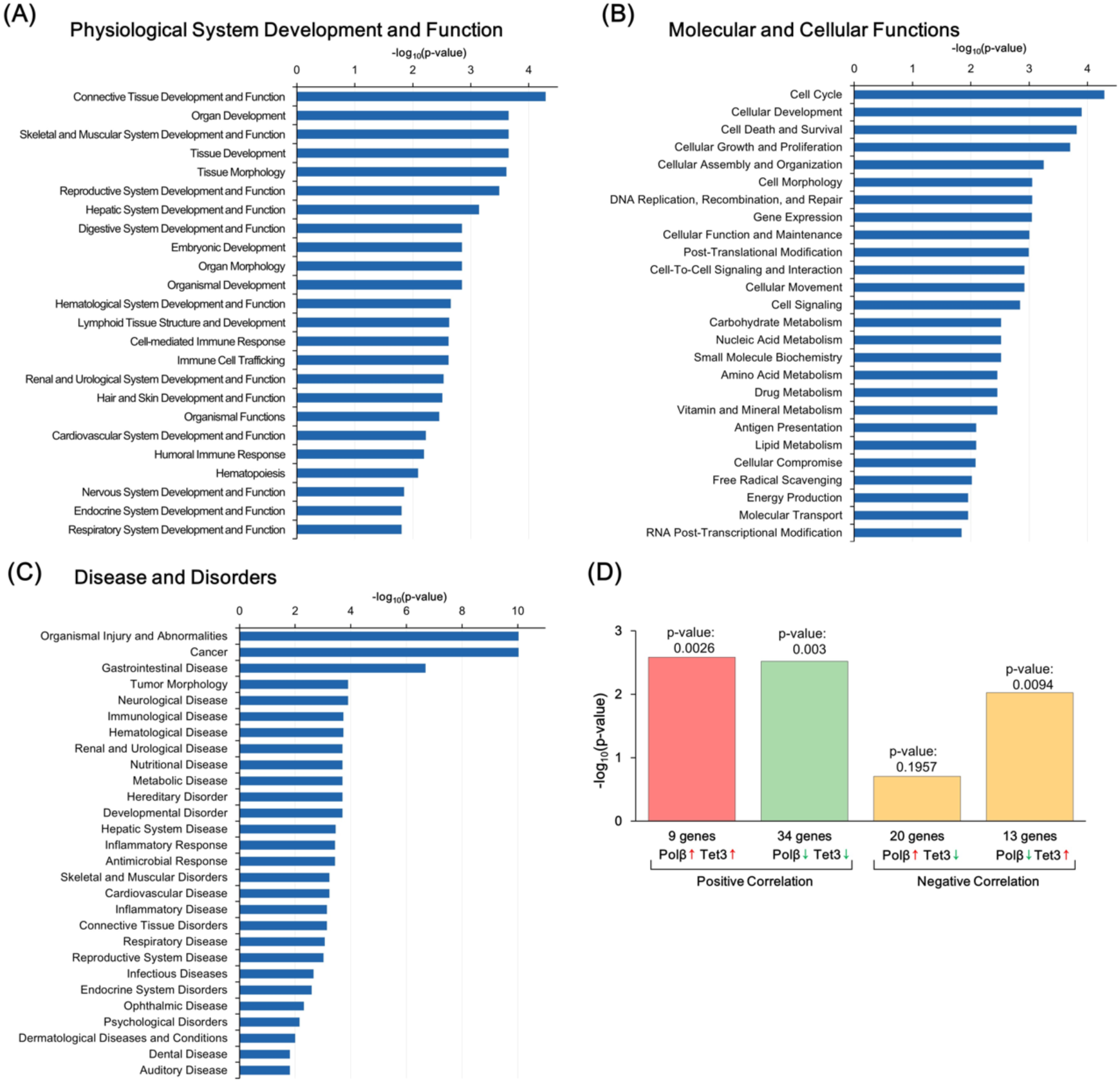
Polβ deficiency affects gene expression and dendrite morphology of hippocampus neurons during postnatal development. (A–C) Functional annotation of DEGs between *Nex-Cre/Polβ^fl/fl^* and *Polβ^fl/fl^* hippocampus in the three primary categories by IPA (p < 0.05, Fisher’s exact test): Physiological System Development and Function (A), Molecular and Cellular Functions (B), and Disease and Disorders (C). (D) Comparative analysis of DEGs between Polβ-deficient hippocampus and TET3 shRNA-transfected hippocampal neuronal culture.

**Supplemental Table 1.** Summary of comprehensive behavioral test battery.

| Test | Measure | p-value |
| --- | --- | --- |
| **General Health** |  |  |
| Body weight | Weight (g) | 0.3496 |
| Body temperature | Temperature (°C) | 0.5477 |
| Grip strength | Strength (N) | 0.8528 |
| Wire hang | Latency to fall (sec) | 0.4578 |
| **Light/dark transition test** |  |  |
|  | Stay time in light (sec) | 0.8748 |
|  | Number of transitions | 0.8044 |
|  | Latency to light (sec) | 0.4936 |
| **Open field test** |  |  |
|  | Total distance (cm) | G: 0.9697, G × T: < 0.0001*** |
|  | Vertical activity | G: 0.4896, G × T: 0.898 |
|  | Center time (sec) | G: 0.8841, G × T: 0.9607 |
|  | Stereotypic counts | G: 0.4577, G × T: 0.0032** |
| **Elevated plus maze test** |  |  |
|  | Entries into open arms (%) | 0.0023** |
|  | Time on open arms (%) | 0.0303* |
| **Rotarod** |  |  |
|  | Latency to fall (sec) | G: 0.0703, G × Tr: 0.825 |
| **Hot plate** |  |  |
|  | Latency (sec) | 0.6464 |
| **Social Interaction (Novel environment)** |  |  |
|  | Total duration of contact (sec) | 0.0445* |
|  | Number of contacts | 0.1277 |
|  | Mean duration / contact (sec) | 0.5133 |
| **3 chamber social approach test** |  |  |
| Sociability | Time spent around stranger cage (%) | 0.7741 |
| Preference | Time spent around stranger cage (%) | 0.0748 |
| **Prepulse inhibition test** |  |  |
| Startle response | Startle amplitude to 110, 120 dB | G: 0.0062, G × S: 0.0421* |
| Prepulse inhibition | PPI (PPI 74, 78 dB - startle 110 dB) (%) | G: 0.1219, G × P: 0.8899 |
|  | PPI (PPI 74, 78 dB - startle 120 dB) (%) | G: 0.0333*, G × P: 0.1127 |
| **Porsolt forced swim** |  |  |
|  | Immobility (%) Day1 | G: 0.1147, G × T: 0.2675 |
|  | Immobility (%) Day2 | G: 0.1134, G × T: 0.0284* |
| **T-maze test (spontaneous alteration)** |  |  |
|  | Correct response (%) | G: 0.0256*, G × Tr: 0.3757 |
|  | Latency (sec) | G: 0.1612, G × Tr: 0.2337 |
| **Barnes maze test** |  |  |
| Acquisition | Number of errors | G: 0.0003***, G × Tr: 0.0031** |
|  | Distance traveled | G: 0.0094**, G × Tr: 0.0058** |
|  | Latency | G: 0.2051, G × Tr: 0.1263 |
|  | Omission errors | G: 0.375, G × Tr: 0.5279 |
| Probe test (1 day) | Time spent around target hole (%) | 0.0031** |
| Probe test (1 month) | Time spent around target hole (%) | 0.0254* |
| **Fear conditioning** |  |  |
| Conditioning | Freezing (%) | G: 0.1434, G × T: 0.883 |
|  | Distance traveled | G: 0.0181*, G × T: 0.5153 |
| Context testing (1 day) | Freezing (%) | G: 0.0809, G × T: 0.8485 |
|  | Distance traveled | G: 0.0195*, G × T: 0.8551 |
| Cued testing with altered context (1 day) | Freezing during CS (%) | G: 0.4288, G × T: 0.8812 |
|  | Distance traveled during CS | G: 0.4428, G × T: 0.9637 |
| Context testing (1 month) | Freezing (%) | G: 0.039*, G × T: 0.2822 |
|  | Total distance traveled | G: 0.0182*, G × T: 0.1719 |
| Cued testing with altered context (1 month) | Freezing during CS (%) | G: 0.1978, G × T: 0.9227 |
|  | Distance traveled during CS | G: 0.2095, G × T: 0.7771 |
| **Tail suspension test** |  |  |
|  | Immobility (%) | G: 0.9557, G × T: 0.1154 |
| **Home cage social interaction** |  |  |
|  | Mean number of particles | G: 0.9939 (light: 0.925, dark: 0.8939) |
|  | Activity level | G: 0.0516 (light: 0.2802, dark: 0.0215*) |

\*p < 0.05, \*\*p < 0.01, and \*\*\*p < 0.001 (n=20 in each group, one-way or two-way repeated measures ANOVA). In two-way repeated measures ANOVA, p-values of interaction of genotype (G) with time (T), trial (Tr), startle (S), or PPI (P) are shown.

